# Epstein–Barr Virus Load in Lymphoblastoid Cell Lines and Its Association with Schizophrenia

**DOI:** 10.64898/2026.11.24.768881

**Authors:** Daniel Yin, Hanwen Zhang, Alan Sanders, Jubao Duan

## Abstract

**Background:** Blood Epstein–Barr virus (EBV) DNAs are reported to have genetic determinants and correlate with prevalence of autoimmune, neurological, respiratory and cardiovascular condition. However, whether a patient is more susceptible to EBV virus infection is unknown. Lymphoblastoid cell lines (LCLs) are often immortalized by infecting individuals’ peripheral lymphocytes by EBV in a dish, providing an ex vivo model to address this question. Here, we examined associations of EBV viral load in LCLs with schizophrenia (SCZ) diagnosis, SCZ and bipolar (BIP) polygenic risk scores, and host genetic variation.

**Methods:** EBV viral load in LCLs from the Molecular Genetics of Schizophrenia (MGS) study was examined using univariable and multivariable linear regression. Separate multivariable models evaluated SCZ diagnosis, bipolar disorder polygenic score (BIP-PGS), and SCZ polygenic score (SCZ-PGS), adjusting for age, sex, and the first five genetic ancestry principal components. An exploratory analysis examined age at schizophrenia onset. A genome-wide association study (GWAS) was conducted in 2,000 participants using the same covariates.

**Results:** SCZ diagnosis was not significantly associated with EBV viral load after covariate adjustment (β = −0.004, SE = 0.025, P = 0.868). Neither BIP-PGS (β = −0.009, SE = 0.013, P = 0.469) nor SCZ-PGS (β = −0.002, SE = 0.013, P = 0.897) was significantly associated with viral load. Univariable analyses also identified no significant association with age at SCZ onset. For GWAS of EBV loads in LCL, no variant reached genome-wide significance (P < 5 × 10^−8^). The strongest association was observed at rs7188098 in the GP2 gene on chromosome 16 (P = 3.7 × 10^−7^).

**Conclusions:** These analyses did not identify statistically significant associations of LCL EBV viral load with SCZ diagnosis or the psychiatric polygenic scores examined. The GWAS identified suggestive signals of association with EBV load, which require further investigation in larger sample sizes.

## 1. Introduction

Epstein–Barr virus (EBV) establishes persistent infection and can transform human B lymphocytes into continuously proliferating lymphoblastoid cell lines (LCLs). These properties have made EBV-transformed LCLs widely used resources for human genetic and molecular research. However, EBV genome abundance varies across LCLs, suggesting that the host and cellular factors may influence this variation and its relevance to disease-associated phenotypes (Houldcroft et al., 2014; Mandage et al., 2017). A recent population-scale study identified host genetic contributions of persistent EBV DNA in blood, however, this phenotype differs from EBV abundance in cultured LCLs (Nyeo et al., 2026). LCLs provide an experimental setting in which to investigate variation in EBV abundance alongside host genotype. Previous studies have examined the genetic architecture of EBV copy number in these cells. Houldcroft et al. (2014) reported evidence of heritability despite identifying no individually significant SNP associations, while Mandage et al. (2017) identified several suggestive genomic regions in LCLs derived from the 1000 Genomes Project. Davies et al. (2010) found that high EBV copy number was accompanied by increased expression of lytic viral genes, indicating that viral abundance can reflect differences in the balance between latency and reactivation. Therefore, whether EBV loads in LCL are associated with any genetic variants with genome-wide significance remains unclear.

Previous research using MGS-derived LCLs demonstrated the utility of this model for investigating differences in transcriptional responses to dopamine between SCZ cases and controls (Sanders et al., 2013; Duan et al., 2018; Kos et al., 2018), as well as gene expression differences under baseline conditions (Sanders et al., 2013, 2017; Duan et al., 2015). Interestingly, a possible relationship between EBV and SCZ has been suggested by studies of antiviral immune responses. Dickerson et al. (2019) reported increased antibody responses to EBV virions and viral capsid antigens in individuals with SCZ, without corresponding increases in antibodies to EBV nuclear antigen-1. These findings suggest that associations may depend on the specific component of the immune response examined. They also highlight the need to distinguish antibody reactivity from measures of viral abundance: an altered immune response does not, by itself, establish increased viral load.

SCZ and bipolar disorder (BIP) have polygenic architectures, with genetic susceptibility distributed across many variants (Trubetskoy et al., 2022; O’Connell et al., 2025). Polygenic scores (PGS) offer an approach to testing whether aggregate genetic liability to these disorders is associated with EBV viral load in LCLs. These analyses will complement case–control comparisons by testing whether aggregate psychiatric genetic liability is associated with the cellular phenotype.

In this study, we investigated EBV viral load in LCLs from the Molecular Genetics of SCZ (MGS) study (Shi et al., 2009; Sanders et al., 2010, 2013; Duan et al., 2018). Using the LCLs that have been assayed for EBV loads in our previous transcriptomic studies (Sanders et al., 2013; Duan et al., 2018), we examined the associations of EBV loads with SCZ diagnosis, BIP disorder and SCZ polygenic scores, and age at SCZ onset. We also conducted a genome-wide association study (GWAS) to identify host genetic variants associated with EBV viral load. Together, these analyses evaluated whether this cellular phenotype was associated with psychiatric diagnosis, aggregate psychiatric genetic liability, or individual host genetic variants.

## 2. Data

### 2.1 EBV and genetic data

Study participants were drawn from the MGS study, which was established to investigate the genetic and molecular factors associated with SCZ. The present study included participants with available EBV viral load measurements, SCZ diagnostic information, demographic characteristics, and genetic ancestry principal components (PCs).

EBV load measurements were obtained from the MGS study LCL dataset. A previous study of MGS samples quantified EBV DNA using TaqMan quantitative real-time PCR (qPCR) on DNA extracted from EBV-transformed LCLs (Sanders et al., 2013). The supplied log-transformed EBV load measurements were used as the quantitative phenotype in the present association analyses. These measurements reflect EBV DNA abundance in cultured LCLs rather than circulating EBV DNA levels or EBV-specific antibody responses in peripheral blood.

Participants were classified as SCZ cases or unaffected controls according to their diagnostic status. Individuals with missing values in variables required for a given regression analysis were excluded from the corresponding analysis.

DNA samples were genotyped using the Affymetrix 6.0 array at the Broad Institute. Genotype quality control procedures have been described previously (Shi et al., 2009). Genotype imputation was performed using IMPUTE2 (version 2.1.2) (Howie et al., 2009, 2011), with the 1000 Genomes Project Phase 3 reference panel (October 2014 release) (Auton et al., 2015). Pre-phasing was conducted using SHAPEIT2 (version v2.r644) to generate best-guess haplotypes (Delaneau et al., 2011). Imputed SNPs were excluded if they had an IMPUTE2 information score below 0.6, an IMPUTE2 certainty score below 0.8, or a minor allele frequency (MAF) below 0.01. Following quality control, 5,132,475 imputed SNPs were retained for subsequent polygenic score analyses.

Of the 2,466 participants with available EBV viral load measurements, 2,002 had genotype data available and were of European ancestry. Two additional participants were excluded because of missing covariate information, resulting in a final analytic sample of 2,000 individuals, including 842 SCZ cases and 1,158 unaffected controls. The mean age was 44.79 years (SD = 11.79), and the sample included 929 females and 1,071 males. Five genetic ancestry PCs (PC1–PC5), calculated from genotype data, were included as covariates in subsequent association analyses.

### 2.2 Polygenic score

Polygenic scores for BIP disorder (BIP-PGS) and SCZ (SCZ-PGS) were calculated for MGS participants using the PRS-CS software (Ge et al., 2019). Two sets of publicly available genome-wide association study summary statistics from the Psychiatric Genomics Consortium were used as discovery datasets.

For BIP disorder, we used the PGC BIP 2024 GWAS summary statistics (bip2024; O’Connell et al., 2025). For SCZ, we used the PGC SCZ Working Group GWAS summary statistics (SCZw3; Trubetskoy et al., 2022).

The corresponding GWAS summary statistics were used as reference inputs for PRS-CS to estimate posterior SNP effect sizes. Individual-level PGS scores were subsequently calculated for MGS participants based on the estimated SNP effect sizes and their genotype data. The resulting scores were used to quantify genetic susceptibility to BIP and SCZ in subsequent association analyses.

## 3. Statistical analysis

Associations of EBV viral load with SCZ diagnosis, SCZ-PGS, and BIP-PGS scores were evaluated using linear regression models in R. Age, sex, and the first five genetic ancestry principal components were included as covariates to account for potential demographic and population stratification effects.

In addition, a GWAS was conducted to identify genetic variants associated with EBV viral load. The analysis included 2,000 participants with available EBV measurements and genotype data. Linear regression models were fitted for each imputed SNP using PLINK, with EBV viral load as the quantitative outcome and the same covariates (age, sex, and PC1–PC5) included in the models.

## 4. Results

### 4.1. Preliminary results

Preliminary univariable linear regression analyses were conducted to examine the associations between EBV viral load and each of the following variables: SCZ, age, sex, PC1, …, PC5, BIP-PGS, SCZ-PGS, and age at SCZ onset. None of these variables were significantly associated with EBV viral load at the 0.05 significance level, as shown in Table 1.

**Table 1:**
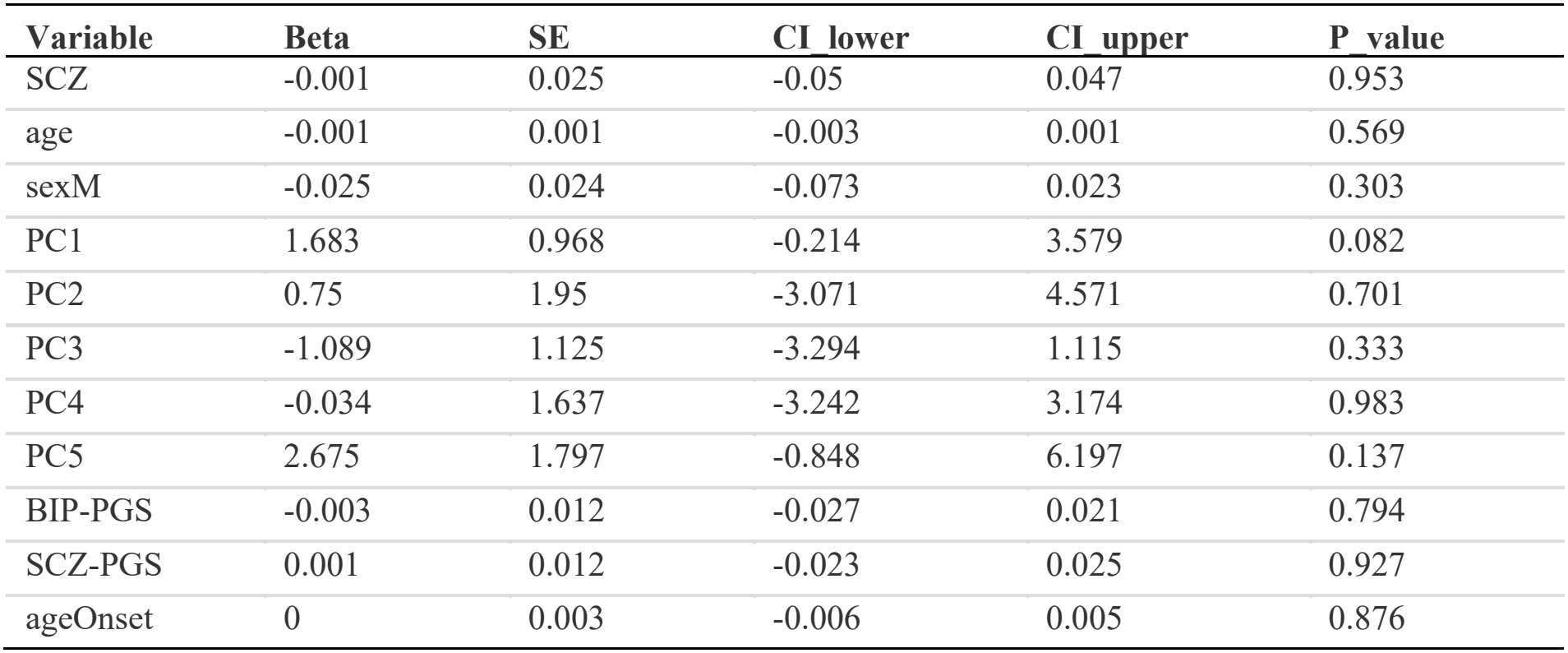
Univariate linear regression analysis of factors associated with EBV viral load.

| **Variable** | **Beta** | **SE** | **CI_lower** | **CI_upper** | **P_value** |
|---|---|---|---|---|---|
| SCZ | -0.001 | 0.025 | -0.05 | 0.047 | 0.953 |
| age | -0.001 | 0.001 | -0.003 | 0.001 | 0.569 |
| sexM | -0.025 | 0.024 | -0.073 | 0.023 | 0.303 |
| PC1 | 1.683 | 0.968 | -0.214 | 3.579 | 0.082 |
| PC2 | 0.75 | 1.95 | -3.071 | 4.571 | 0.701 |
| PC3 | -1.089 | 1.125 | -3.294 | 1.115 | 0.333 |
| PC4 | -0.034 | 1.637 | -3.242 | 3.174 | 0.983 |
| PC5 | 2.675 | 1.797 | -0.848 | 6.197 | 0.137 |
| BIP-PGS | -0.003 | 0.012 | -0.027 | 0.021 | 0.794 |
| SCZ-PGS | 0.001 | 0.012 | -0.023 | 0.025 | 0.927 |
| ageOnset | 0 | 0.003 | -0.006 | 0.005 | 0.876 |

Figure 1 presents the distribution of EBV viral load among individuals with SCZ and healthy controls. The boxplots showed substantial overlap in the distributions of EBV viral load between the two groups, with no apparent difference in the median viral load or overall distribution. Figure 2 illustrates the relationship between EBV viral load and age at onset of SCZ. The scatterplot showed considerable variability in EBV viral load across the range of age at onset, with no clear increasing or decreasing trend. The fitted regression line was relatively flat, and the 95% confidence interval indicated uncertainty around the estimated relationship. Overall, the scatterplot provided no apparent evidence of a linear association between EBV viral load and age at onset of SCZ.

**Figure 1.**
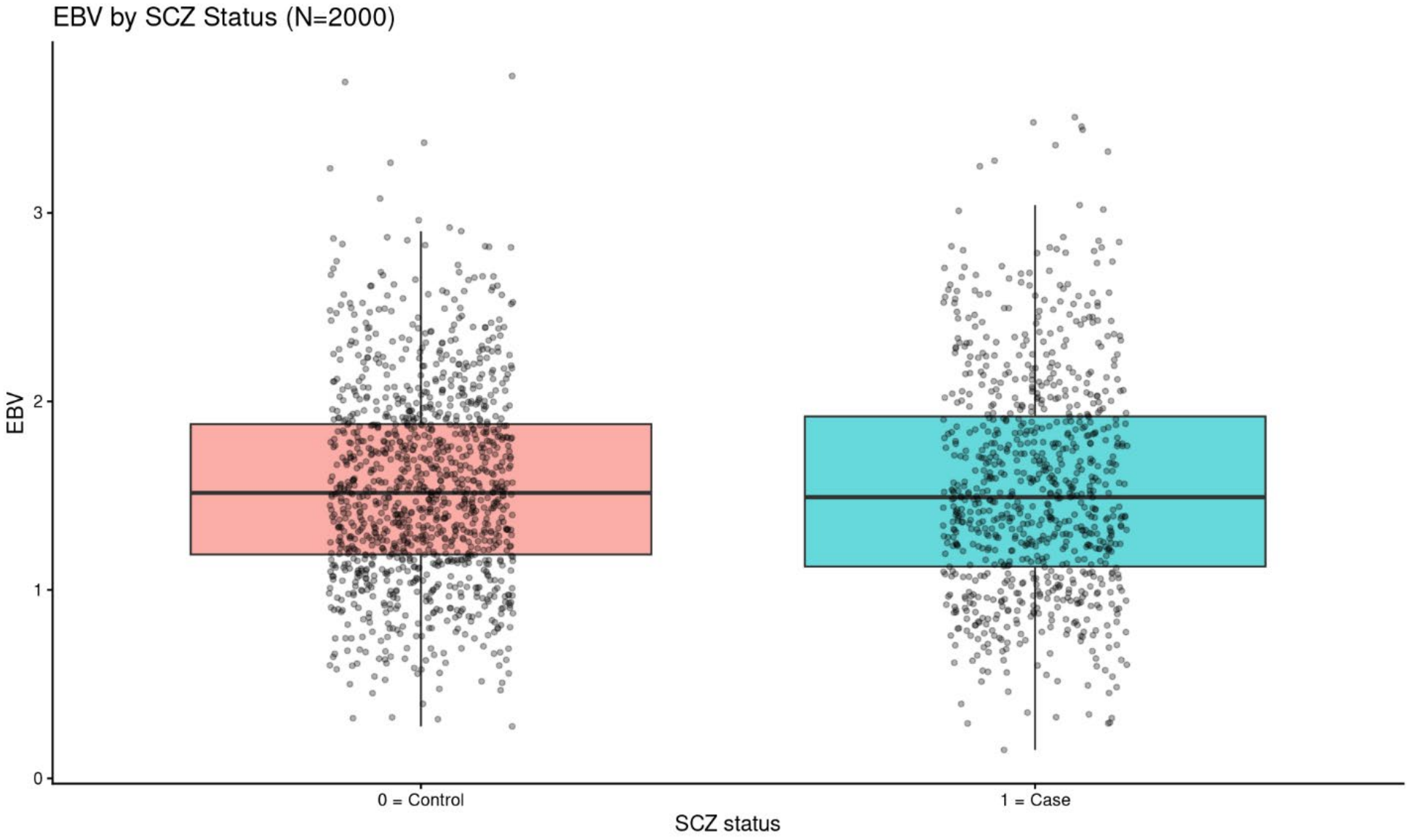
Boxplots of EBV viral load between SCZ cases (N=842) and controls (N=1158).

**Figure 2.**
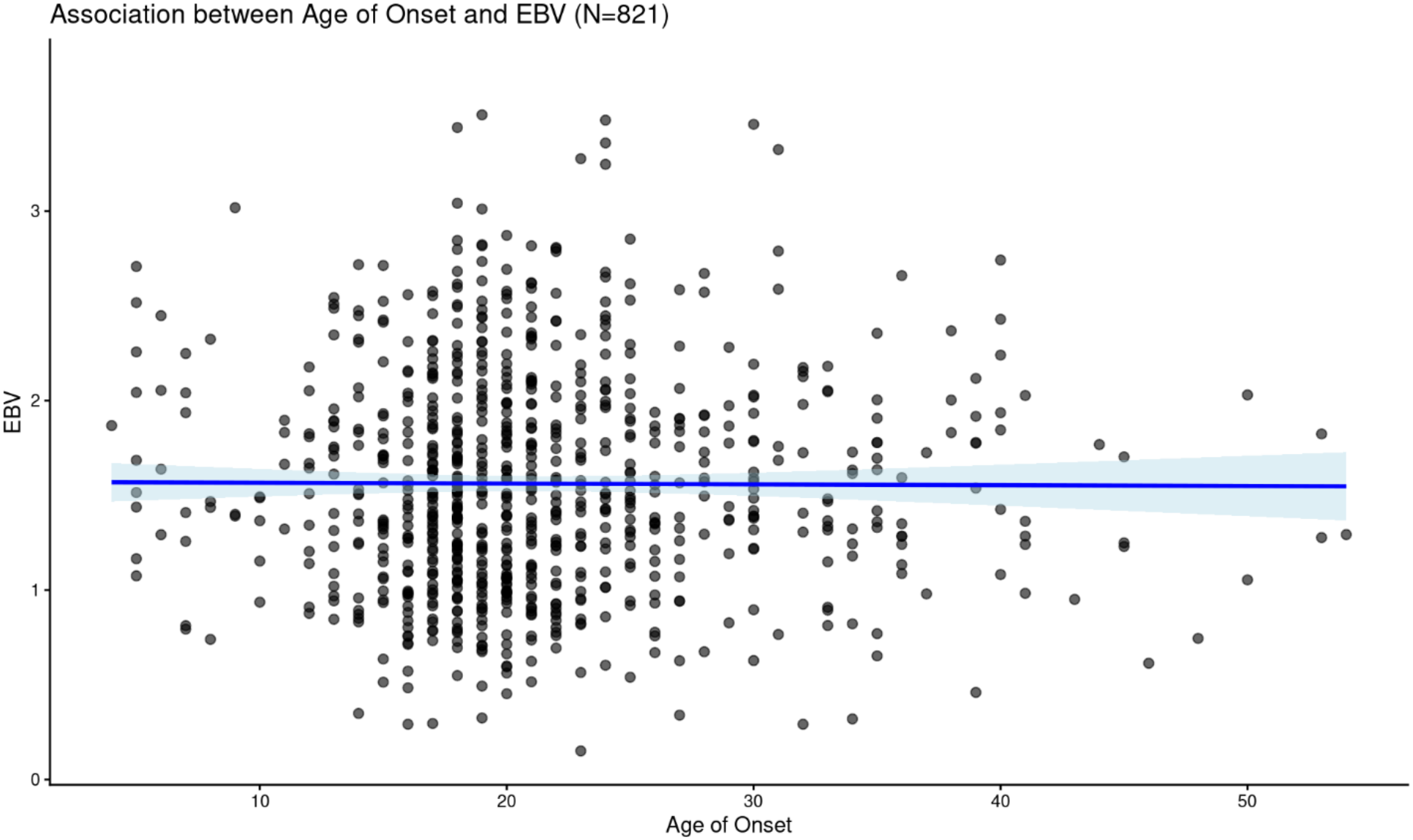
Scatterplot showing the association between EBV viral load and age at onset of SCZ. The solid line represents the fitted linear regression line, and the shaded area indicates the 95% confidence interval for the regression line.

### 4.2. Association between SCZ diagnosis and EBV viral load

To further investigate the association between SCZ diagnosis and EBV viral load, a multivariable linear regression analysis was performed, adjusting for age, sex, and the first five genetic ancestry PCs. As shown in Table 2, SCZ diagnosis was not significantly associated with EBV viral load after adjustment for age, sex, and genetic ancestry principal components (β = −0.004, SE = 0.025, 95% CI: −0.053 to 0.045, p = 0.868). These findings suggest that EBV viral load measured in MGS lymphoblastoid cell lines was not significantly different between SCZ cases and unaffected controls after accounting for the demographic and genetic ancestry covariates.

**Table 2:**
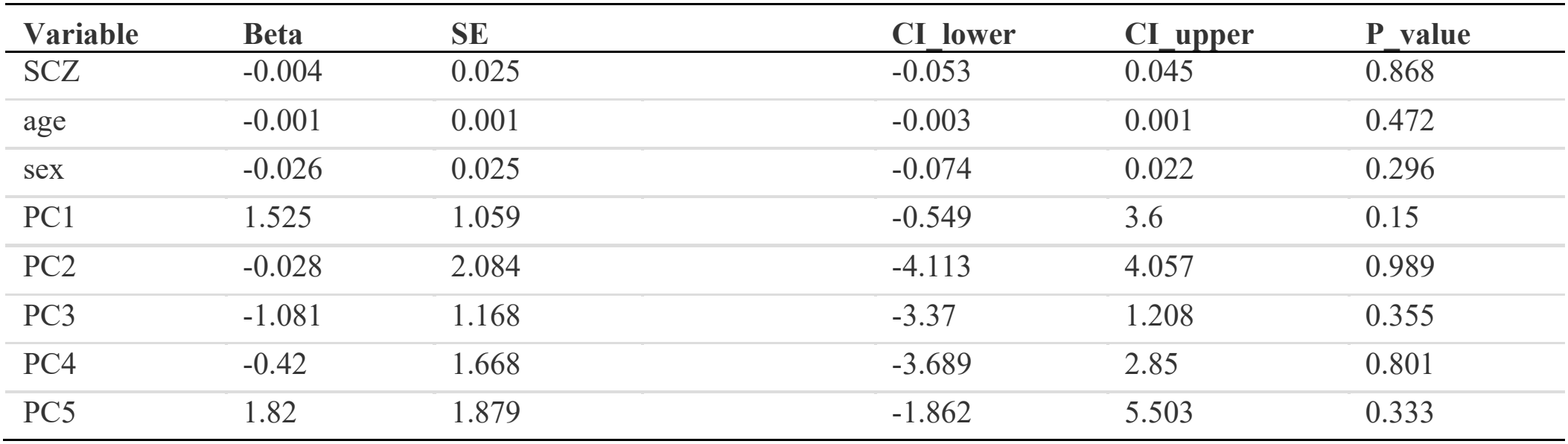
Multivariable linear regression analysis of the association between SCZ diagnosis and EBV viral load, adjusted for age, sex, and genetic ancestry principal components (PC1–PC5).

| **Variable** | **Beta** | **SE** | **CI_lower** | **CI_upper** | **P_value** |
|---|---|---|---|---|---|
| SCZ | -0.004 | 0.025 | -0.053 | 0.045 | 0.868 |
| age | -0.001 | 0.001 | -0.003 | 0.001 | 0.472 |
| sex | -0.026 | 0.025 | -0.074 | 0.022 | 0.296 |
| PC1 | 1.525 | 1.059 | -0.549 | 3.6 | 0.15 |
| PC2 | -0.028 | 2.084 | -4.113 | 4.057 | 0.989 |
| PC3 | -1.081 | 1.168 | -3.37 | 1.208 | 0.355 |
| PC4 | -0.42 | 1.668 | -3.689 | 2.85 | 0.801 |
| PC5 | 1.82 | 1.879 | -1.862 | 5.503 | 0.333 |

### 4.3. Association of BIP-PGS and SCZ-PGS scores with EBV viral load

To further investigate the associations of genetic predisposition to BIP and SCZ with EBV viral load, separate multivariable linear regression analyses were performed using the BIP-PGS and SCZ-PGS as predictors. Both models were adjusted for age, sex, and the first five genetic ancestry PCs.

As shown in Table 3, neither BIP-PGS nor SCZ-PGS was significantly associated with EBV viral load after adjustment for the covariates. Specifically, BIP-PGS showed no significant association with EBV viral load (β = −0.009, SE = 0.013, p = 0.469). Similarly, SCZ-PGS was not significantly associated with EBV viral load (β = −0.002, SE = 0.013, p = 0.897).

**Table 3:** Associations of bipolar disorder and SCZ polygenic scores with EBV viral load, adjusted for age, sex, and genetic ancestry principal components.

|  | **BIP-PGS** |  |  | **SCZ-PGS** |  |  |
|---|---|---|---|---|---|---|
| **Variable** | **Beta** | **SE** | **P_value** | **Beta** | **SE** | **P_value** |
| *PGS* | *-0.009* | *0.013* | *0.469* | *-0.002* | *0.013* | *0.897* |
| age | -0.001 | 0.001 | 0.474 | -0.001 | 0.001 | 0.47 |
| sex | -0.026 | 0.024 | 0.296 | -0.026 | 0.024 | 0.294 |
| PC1 | 1.703 | 1.086 | 0.117 | 1.531 | 1.072 | 0.153 |
| PC2 | -0.037 | 2.083 | 0.986 | -0.038 | 2.089 | 0.986 |
| PC3 | -1.051 | 1.167 | 0.368 | -1.086 | 1.171 | 0.354 |
| PC4 | -0.46 | 1.667 | 0.783 | -0.423 | 1.671 | 0.8 |
| PC5 | 1.846 | 1.877 | 0.325 | 1.812 | 1.885 | 0.337 |

These findings suggest that neither BIP-PGS nor SCZ-PGS scores were significantly associated with EBV viral load in this sample, indicating no evidence of an association between genetic predisposition to these psychiatric disorders and EBV viral load.

### 4.4. GWAS analysis of EBV Viral Load

A GWAS of EBV viral load was conducted in 2,000 participants, adjusting for age, sex, and the first five genetic ancestry PCs. The Manhattan plot in Figure 3 illustrates the distribution of association signals across the genome. Among variants with a minor allele frequency greater than 0.01, the strongest association was observed at rs7188098 on chromosome 16 (β = 0.19, P = 3.67 × 10^−7^), with *GP2 (glycoprotein 2)* identified as the nearest gene. This was followed by rs139670902 on chromosome 5 (β = 0.15, P = 1.61 × 10^−6^), nearest to *ADAMTS2*; rs116566441 on chromosome 2 (β = −0.29, P = 2.07 × 10^−6^), nearest to *NPPC*; and rs13393575 on chromosome 2 (β = −0.11, P = 2.09 × 10^−6^), nearest to *GYPC*.

**Figure 3.**
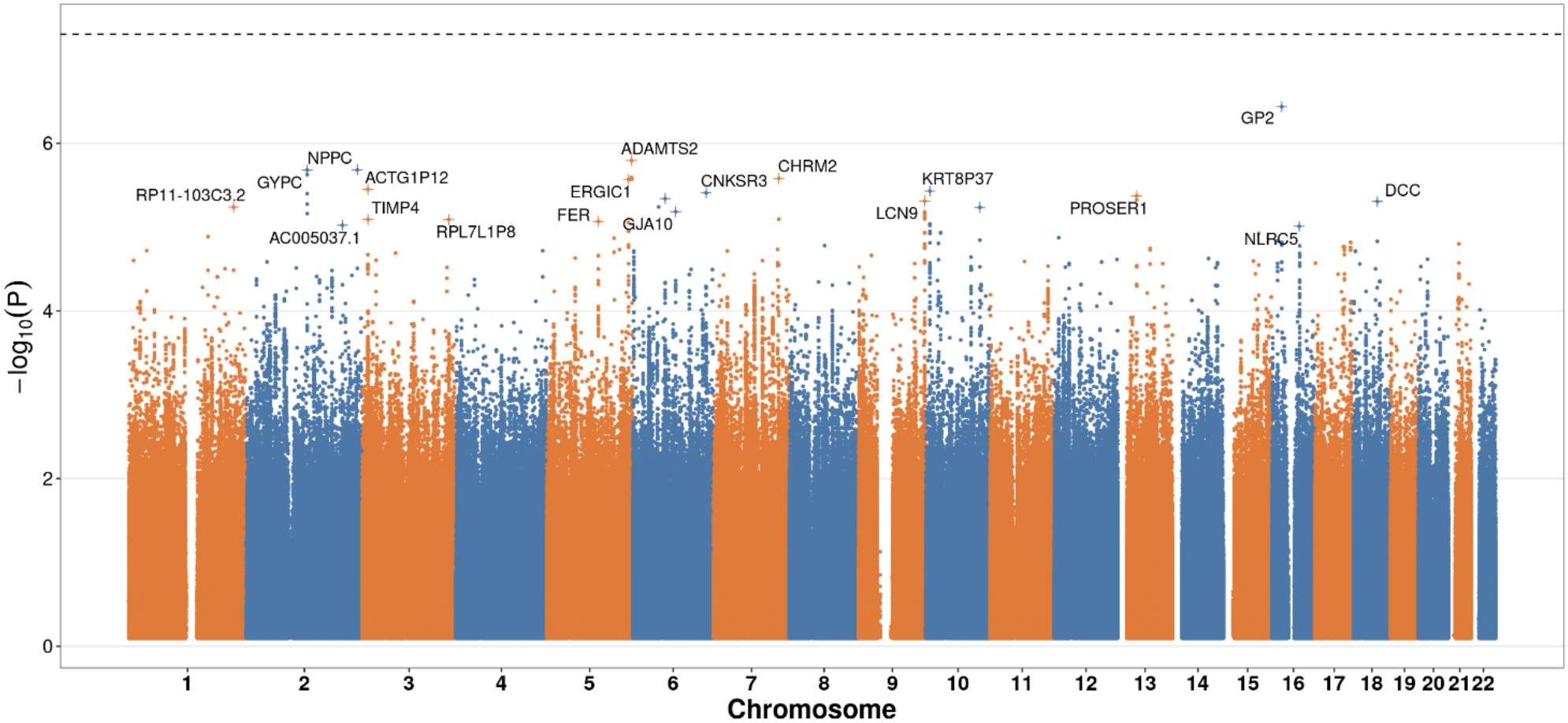
Manhattan plot of genome-wide association results for EBV viral load in 2,000 participants. Association analyses were performed using linear regression in PLINK, adjusting for age, sex, and the first five genetic ancestry principal components. The x-axis represents the genomic position of each SNP by chromosome, and the y-axis represents the −log_10_(P-value). Gene annotation was performed using a 100-kb window around each SNP, and the nearest gene was highlighted for SNPs with P < 1 × 10^−5^. The horizontal dashed line indicates the genome-wide significance threshold (P < 5 × 10^−8^).

Although several variants showed suggestive evidence of association with EBV viral load, none reached the genome-wide significance threshold of P < 5 × 10^−8^. The quantile–quantile plot is presented in Figure 4. The genomic inflation factor (λ_GC_ = 1.002) indicated minimal overall inflation of the association test statistics. Additional top association signals with P < 1 × 10^−5^ are presented in Table 4. Gene annotations, including the nearest genes and genes located within a 100-kb window of each variant, were obtained using the Ensembl Variant Effect Predictor (VEP; https://www.ensembl.org/tools/vep).

**Table 4:** Top SNPs associated with EBV viral load in the GWAS, including genomic positions, allele frequencies, effect estimates, P values, and nearest genes identified within a 100-kb window.

| SNP | CHR | BP | A1 | MAF | BETA | STAT | P | GENE | DISTANCE (bp) |
|---|---|---|---|---|---|---|---|---|---|
| rs7188098 | 16 | 20326407 | T | 0.060 | 0.193 | 5.103 | 3.67E-07 | GP2 | 3317 |
| rs139670902 | 5 | 178528300 | C | 0.107 | 0.147 | 4.816 | 1.61E-06 | ADAMTS2 | 9552 |
| rs116566441 | 2 | 232882777 | C | 0.022 | -0.287 | -4.761 | 2.07E-06 | NPPC | 91664 |
| rs13393575 | 2 | 127477191 | T | 0.293 | -0.108 | -4.763 | 2.09E-06 | GYPC | 22945 |
| rs324667 | 7 | 136728372 | C | 0.367 | -0.085 | -4.712 | 2.63E-06 | CHRM2 | 23370 |
| rs9637855 | 5 | 172306540 | T | 0.238 | 0.101 | 4.708 | 2.70E-06 | ERGIC1 | 25680 |
| rs307582 | 3 | 12085149 | T | 0.192 | 0.100 | 4.649 | 3.55E-06 | ACTG1P12 | 26546 |
| rs10795656 | 10 | 8595839 | A | 0.279 | -0.092 | -4.641 | 3.70E-06 | KRT8P37 | 38770 |
| rs62432754 | 6 | 154846757 | A | 0.264 | -0.128 | -4.644 | 3.90E-06 | CNKSR3 | 14964 |
| rs9315642 | 13 | 39547702 | T | 0.362 | 0.119 | 4.624 | 4.26E-06 | PROSER1 | 36301 |
| rs9454303 | 6 | 68536629 | G | 0.150 | 0.112 | 4.597 | 4.57E-06 | intergenic |  |
| rs484198 | 9 | 138558241 | A | 0.070 | 0.158 | 4.582 | 4.90E-06 | LCN9 | 292 |
| rs9964756 | 18 | 50441817 | T | 0.113 | 0.123 | 4.580 | 4.95E-06 | DCC | 8337 |
| rs112829384 | 1 | 221718413 | A | 0.011 | 0.385 | 4.547 | 5.77E-06 | RP11-103C3.2 | 4715 |
| rs2792728 | 10 | 113774442 | T | 0.205 | -0.104 | -4.547 | 5.81E-06 | intergenic |  |
| rs2236181 | 6 | 90661576 | C | 0.319 | 0.090 | 4.522 | 6.55E-06 | GJA10 | 55737 |
| rs61289654 | 3 | 12190384 | T | 0.174 | -0.108 | -4.476 | 8.10E-06 | TIMP4 | 4167 |
| rs67460045 | 3 | 182321031 | A | 0.092 | -0.191 | -4.485 | 8.13E-06 | RPL7L1P8 | 7431 |
| rs7717569 | 5 | 108505337 | C | 0.119 | 0.135 | 4.466 | 8.56E-06 | FER | 7881 |
| rs6743474 | 2 | 201706901 | T | 0.199 | -0.126 | -4.449 | 9.51E-06 | AC005037.1 | 2062 |
| rs42221 | 16 | 57113229 | A | 0.192 | 0.105 | 4.436 | 9.77E-06 | NLRC5 | 1502 |

**Figure 4.**
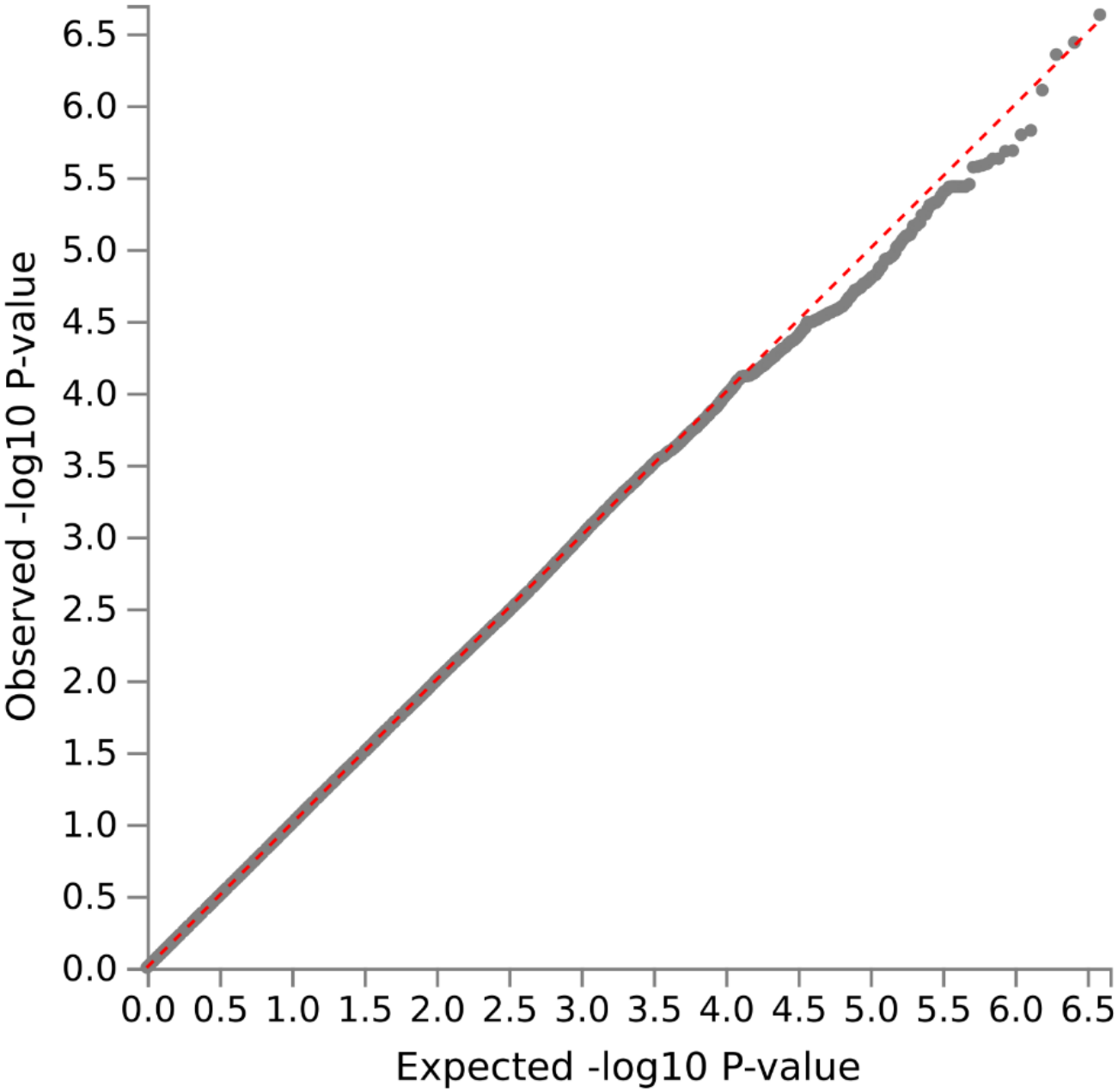
Quantile–quantile plot of the genome-wide association analysis of EBV viral load.

Among the genes identified near the top SNP-associated loci, several have biological functions potentially relevant to immune regulation and SCZ. *NLRC5* is particularly noteworthy because of its established role in regulating major histocompatibility complex (MHC) class I gene expression and antigen presentation, which are essential for *CD8+* T-cell-mediated antiviral immune responses (Meissner et al., 2010; Neerincx et al., 2010). Given the importance of cellular immunity in controlling persistent viral infections, *NLRC5* represents a biologically plausible candidate for further investigation in relation to EBV viral load. *GP2*, which was located near the strongest association signal (rs7188098), is another interesting candidate because of its reported role as a receptor mediating bacterial antigen uptake by intestinal M cells and initiating mucosal immune responses (Hase et al., 2009). Although these findings do not establish a direct role for *GP2* in EBV infection, they suggest a potential connection between host immune regulation and variation in viral load. In addition, *ADAMTS2* has an established role in extracellular matrix organization through collagen processing and has shown altered expression in peripheral blood mononuclear cells from patients with SCZ (Vanlerberghe et al., 2025; Kos et al., 2018). Other genes, including *DCC* and *CHRM2*, have previously been investigated in the context of SCZ, particularly in relation to neurodevelopment, neuronal connectivity, and autonomic function (Grant et al., 2012; Wang et al., 2018; Refisch et al., 2023). Collectively, these candidate genes suggest several potentially relevant biological pathways, including antiviral antigen presentation, mucosal immunity, extracellular matrix organization, and neurodevelopment. However, further functional studies are needed to determine whether the associated variants influence the expression or function of these nearby genes and whether these pathways contribute to interindividual variation in EBV viral load.

## 5. Discussion

This study examined EBV viral load in MGS-derived LCLs using clinical association analyses, psychiatric polygenic scores, and a GWAS. SCZ diagnosis was not significantly associated with EBV viral load after adjustment for age, sex, and genetic ancestry PCs. Neither BIP-PGS nor SCZ-PGS scores showed a significant association, and preliminary analyses found no significant association with age at SCZ onset.

The adjusted effect of SCZ diagnosis was close to zero, but its confidence interval included both negative and positive effects. This finding therefore does not establish equivalence between cases and controls. Similarly, the PGS results suggest that the genetic liability captured by these scores did not detectably predict LCL EBV viral load in this sample. They do not exclude small effects or associations with other aspects of EBV biology.

No variant reached genome-wide significance in the GWAS of EBV viral load. The strongest signal was observed at rs7188098 near *GP2* on chromosome 16, followed by rs139670902 near *ADAMTS2* on chromosome 5. *GP2* has been implicated in antigen uptake and mucosal immune responses (Hase et al., 2009), while *ADAMTS2* is involved in extracellular matrix organization and has been reported to show altered expression in peripheral blood mononuclear cells from patients with SCZ (Kos et al., 2018; Vanlerberghe et al., 2025). However, the relevance of these genes to EBV viral load remains uncertain, and these signals should be considered exploratory pending independent replication.

Several limitations should be considered. First, EBV viral load in transformed LCLs may not reflect viral activity or immune control in the body. Second, culture and laboratory variables were not included in the reported models; differences in transformation batch, passage history, or assay conditions could contribute to variation. Third, the GWAS sample size and the MAF of several leading variants limit the certainty of these signals. Finally, the study cannot establish causal relationships, largely due to lack of study power.

Overall, we found no statistically significant associations between EBV viral load in MGS-derived LCLs and SCZ diagnosis, the psychiatric PGS examined, or age at SCZ onset. Larger studies with any resultant significant findings followed up by independent replication, assessment of laboratory factors, and complementary EBV measurements are needed to clarify these relationships.

## Acknowledgement

We thank the study participants of MGS. MGS was mainly supported by R01MH059571, R01MH081800, and U01MH079469 (to P.V.G.); and other NIH grants for other MGS sites (R01MH067257 to N.G.B., R01MH059588 to B.J.M., R01MH059565 to R.F., R01MH059587 to F.A., R01MH060870 to W.F.B., R01MH059566 to D.W.B., R01MH059586 to J.M.S., R01MH061675 to D.F.L., R01MH060879 to C.R.C., U01MH046276 to C.R.C., and U01MH079470 to D.F.L). The transcriptomic work was primarily supported by RC2MH090030 (to A.R.S.), R01MH094116 (to H.H.H.G.), and R01MH094091 (to P.V.G.). Part of this work was conducted in facilities constructed under the support of NIH grant 1C06RR020547. This work was also supported by Endeavor Health Research Institute Scientific Advisory Group (SAG) awards to A.R.S and J.D..

## References

1. Auton A, Abecasis GR, Altshuler DM, et al. A global reference for human genetic variation. Nature. 2015;526:68–74. doi: 10.1038/nature15393.

2. Davies ML, Xu S, Lyons-Weiler J, et al. Cellular factors associated with latency and spontaneous Epstein-Barr virus reactivation in B-lymphoblastoid cell lines. Virology. 2010;400(1):53–67. doi: 10.1016/j.virol.2010.01.002.

3. Delaneau O, Zagury JF, Marchini J. Improved whole-chromosome phasing for disease and population genetic studies. Nat Methods. 2013;10(1):5–6. doi: 10.1038/nmeth.2307.

4. Dickerson F, Jones-Brando L, Ford G, et al. Schizophrenia is associated with an aberrant immune response to Epstein-Barr virus. Schizophr Bull. 2019;45(5):1112–1119. doi: 10.1093/schbul/sby164.

5. Duan J, Sanders AR, Moy W, et al. Transcriptome outlier analysis implicates schizophrenia susceptibility genes and enriches putatively functional rare genetic variants. Hum Mol Genet. 2015;24(16):4674–4685. doi: 10.1093/hmg/ddv199.

6. Duan J, Göring HHH, Sanders AR, et al. Transcriptomic signatures of schizophrenia revealed by dopamine perturbation in an ex vivo model. Transl Psychiatry. 2018;8(1):158. doi: 10.1038/s41398-018-0216-5. PMID: 30115913.

7. Ge T, Chen CY, Ni Y, et al. Polygenic prediction via Bayesian regression and continuous shrinkage priors. Nat Commun. 2019;10:1776. doi: 10.1038/s41467-019-09718-5.

8. Grant A, Fathalli F, Rouleau G, et al. Association between schizophrenia and genetic variation in DCC: a case-control study. Schizophr Res. 2012;137(1–3):26–31. doi: 10.1016/j.schres.2012.02.023.

9. Hase K, Kawano K, Nochi T, et al. Uptake through glycoprotein 2 of FimH+ bacteria by M cells initiates mucosal immune response. Nature. 2009;462(7270):226–230. doi: 10.1038/nature08529.

10. Houldcroft CJ, Petrova V, Liu JZ, et al. Host genetic variants and gene expression patterns associated with Epstein-Barr virus copy number in lymphoblastoid cell lines. PLoS One. 2014;9(10):e108384. doi: 10.1371/journal.pone.0108384.

11. Howie BN, Donnelly P, Marchini J. A flexible and accurate genotype imputation method for the next generation of genome-wide association studies. PLoS Genet. 2009;5(6):e1000529. doi: 10.1371/journal.pgen.1000529.

12. Howie B, Marchini J, Stephens M. Genotype imputation with thousands of genomes. G3 (Bethesda). 2011;1(6):457–470. doi: 10.1534/g3.111.001198.

13. Kos MZ, Duan J, Sanders AR, et al. Dopamine perturbation of gene co-expression networks reveals differential response in schizophrenia for translational machinery. Transl Psychiatry. 2018;8(1):278. doi: 10.1038/s41398-018-0325-1. PMID: 30546022.

14. Mandage R, Telford M, Rodríguez JA, et al. Genetic factors affecting EBV copy number in lymphoblastoid cell lines derived from the 1000 Genome Project samples. PLoS One. 2017;12(6):e0179446. doi: 10.1371/journal.pone.0179446.

15. Meissner TB, Li A, Biswas A, et al. NLR family member NLRC5 is a transcriptional regulator of MHC class I genes. Proc Natl Acad Sci USA. 2010;107(31):13794–13799. doi: 10.1073/pnas.1008684107.

16. Neerincx A, Lautz K, Menning M, et al. A role for the human nucleotide-binding domain, leucine-rich repeat-containing family member NLRC5 in antiviral responses. J Biol Chem. 2010;285(34):26223–26232. doi: 10.1074/jbc.M110.109736.

17. Nyeo SS, Cumming EM, Burren OS, et al. Population-scale sequencing resolves determinants of persistent EBV DNA. Nature. 2026;650:664–672. doi: 10.1038/s41586-025-10020-2.

18. O’Connell KS, Koromina M, van der Veen T, et al. Genomics yields biological and phenotypic insights into bipolar disorder. Nature. 2025;639:968–975. doi: 10.1038/s41586-024-08468-9.

19. Refisch A, Komatsuzaki S, Ungelenk M, et al. Associations of common genetic risk variants of the muscarinic acetylcholine receptor M2 with cardiac autonomic dysfunction in patients with schizophrenia. World J Biol Psychiatry. 2023;24(1):1–11. doi: 10.1080/15622975.2022.2043561.

20. Sanders AR, Drigalenko EI, Duan J, et al. Transcriptome sequencing study implicates immune-related genes differentially expressed in schizophrenia: new data and a meta-analysis. Transl Psychiatry. 2017;7(4):e1093. doi: 10.1038/tp.2017.47.

21. Sanders AR, Göring HHH, Duan J, et al. Transcriptome study of differential expression in schizophrenia. Hum Mol Genet. 2013;22(24):5001–5014. doi: 10.1093/hmg/ddt350.

22. Sanders AR, Levinson DF, Duan J, et al. The Internet-based MGS2 control sample: self report of mental illness. Am J Psychiatry. 2010;167(7):854–865. doi: 10.1176/appi.ajp.2010.09071050.

23. Shi J, Levinson DF, Duan J, et al. Common variants on chromosome 6p22.1 are associated with schizophrenia. Nature. 2009;460:753–757. doi: 10.1038/nature08192.

24. Trubetskoy V, Pardiñas AF, Qi T, et al. Mapping genomic loci implicates genes and synaptic biology in schizophrenia. Nature. 2022;604:502–508. doi:10.1038/s41586-022-04434-5.

25. Vanlerberghe R, Colige A, Malfait AM, et al. ADAMTS2: More than a procollagen N-proteinase. Genes Dis. 2025;12(6):101686. doi: 10.1016/j.gendis.2025.101686.

26. Wang Z, Li P, Wu T, et al. Axon guidance pathway genes are associated with schizophrenia risk. Exp Ther Med. 2018;16(6):4519–4526. doi: 10.3892/etm.2018.6781.

